# Development of pre-clinical murine models for fibrolamellar hepatocellular carcinoma

**DOI:** 10.1101/2023.12.06.569624

**Authors:** Andy He, Garry L. Coles, Griffin G. Hartmann, Julia Arand, Vicky Le, Brandon Mauch, Lei Xu, Thuyen Nguyen, Florette K. Hazard, Julien Sage

## Abstract

Fibrolamellar hepatocellular carcinoma (FLC) is a rare form of cancer that affects primarily adolescents and young adults. FLC tumors are typically associated with an intrachromosomal deletion resulting in expression of a fusion protein between the chaperone DNAJ1B and the protein kinase PKA. FLC is challenging to study because of its rarity and limited pre-clinical models. Here we developed a novel transgenic mouse model of FLC. In this model, DNAJ1B-PKA expression in the liver of mouse embryos results in perinatal lethality associated with liver developmental defects, while DNAJ1B-PKA expression in the liver of adult mice initiates tumors resembling FLC at low penetrance. We sought to develop *ex vivo* cell models from these tumors but failed to establish long-term cell lines. New pre-clinical models of FLC will provide novel insights into the biology of this rare cancer and may help identify novel therapeutic strategies.

## INTRODUCTION

While hepatocellular carcinoma (HCC) in adults is a leading cause of mortality worldwide [1], liver cancer is much rarer in children and young adults. A majority of pediatric liver cancer cases are hepatoblastomas but primary HCCs are sometimes diagnosed [2]. In particular, fibrolamellar hepatocellular carcinoma (FL-HCC, or FLC) is a very rare and distinct subtype of HCC that affects adolescents and young adults. FLC develops in the setting of normal healthy livers, with no known genetic predisposition factor [3-5]. FLC is associated with better survival than HCC in adults, presumably due to the young age of the patients and the lack of cirrhosis, which makes more aggressive surgery possible [6]. However, few therapeutic options have been successfully explored beyond surgery. The 5-year survival rate ranges from 35-75% in patients treated with liver transplantation or resection. In cases where surgery is not possible, the survival drops to 12-14 months [7-10].

Morphologically, FLC tumors are characterized by pleomorphic malignant hepatocytes with large central nucleoli and abundant eosinophilic cytoplasm. The cancer cells are arranged in variably sized nests or cords set within a meshwork of lamellated collagen fibers. In the majority of cases, FLC cells contain hyaline droplets and some isolated cells have round cytoplasmic inclusions known as “pale bodies” [11, 12]. FLC express markers associated with biliary (e.g., CK7 and MUC1) and hepatocytic (e.g., Heppar-1, Albumin, α-1-antitrypsin, and Glypican-3) differentiation, as well as hepatic progenitor/stem cells (e.g., CK19, EPCAM) [3, 13-19].

At the genetic level, a breakthrough in the field was the discovery of a fusion event resulting in high levels of the catalytic subunit of PKA (protein kinase A) in FLC tumors [20, 21]. This finding was confirmed in other studies [22-24]. Rare cases of FLC have been associated with genetic inactivation of *PRKAR1A*, which encodes a regulatory subunit of PKA [25]. Rare cases of *BAP1* mutant HCC also have features of FLC and show amplification of the *PRKACA* gene coding for PKA with no fusion event [26]. Thus, high PKA activity may be sufficient to drive FLC development. Indeed, expression of PKA in the liver of young adult mice is sufficient to initiate tumors resembling FLC [27, 28]. In addition, inhibition of PKA kinase activity has anti-tumor activity against pre-clinical models of FLC [29] and knock-down of the fusion protein inhibits the growth of patient-derived xenografts [30]. However, the small part of DNAJ1B fused to PKA in FLC tumors also contributes to tumor development in mice [27], an observation supported by biochemical and structural studies [31-36]. Few additional alterations have been found in the genome of FLC tumors, but accumulating evidence suggests a role for telomerase activity, MAPK signaling, and WNT signaling [26, 27, 35, 37-41].

The very low incidence rate of 0.02 per 100,000 has kept FLC an understudied cancer [42]. Very few cell lines and patient-derived xenograft models remain available, and these models often do not propagate easily, even though they can provide important model systems [16, 43, 44]. Similarly, mouse models have also been developed using CRISPR/Cas9 approaches; in these models, tumors recapitulate some key features of human FLC, but they grow slowly, limiting their use [27, 28]. Organoids derived from tumors have only been recently described [45, 46]. Here we engineered a new mouse allele in which the DNAJ1B-PKA fusion can be induced following Cre-mediated recombination in transgenic mice. We used this new model to investigate the consequences of DNAJ1B-PKA expression in the adult liver. We were able to develop allografts and 3D culture systems that are amenable to investigations exploring further the biology of FLC and identifying new therapeutic approaches.

## RESULTS

### Transgenic mice expressing DNAJ1B-PKA

We generated transgenic mice by inserting a *DNAJB1-PRKACA-GFP* cDNA into the *Rosa26* locus. Upon Cre-mediated recombination and self-cleaving of the fusion protein, this construct allows expression of the DNAJB1-PKA fusion found in human FLC and the GFP reporter (**Fig. 1A**). *Rosa26*^*LSL-DNAJB1-PKA*^ transgenic mice were healthy and fertile, as would be expected for a conditional allele.

**Figure 1.**
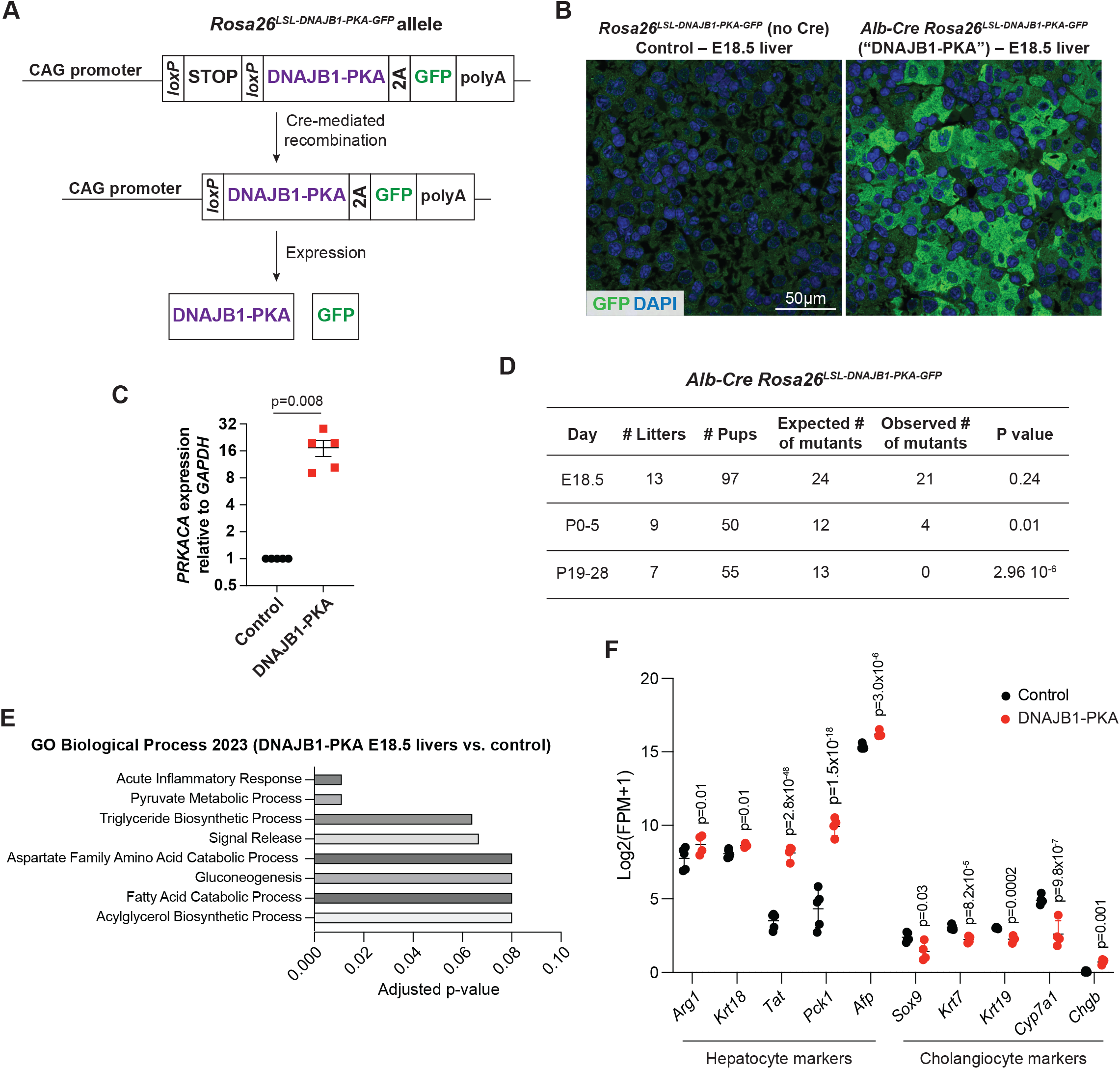
Generation of a transgenic mouse model to express the DNAJ1B-PRAKA fusion. **A**. Schematic representation of the transgene inserted into the *Rosa26* locus. The fusion cDNA is under the control of the CMV early enhancer/chicken β actin (CAG) promoter but is only expressed after Cre-mediated recombination of a transcriptional STOP cassette flanked by *loxP* sites. Upon translation, a self-cleaving 2A peptide separates the DNAJB1-PKA fusion protein from the GFP reporter. **B**. Representative confocal immunofluorescence images for GFP expression (green) from sections of control mice (transgenic mice with no Cre expression) or mice expression the fusion protein and GFP at E18.5. DAPI shows DNA staining (blue). Scale bar: 50µm. **C**. Expression levels of *PRKCA* transcripts analyzed by RT-qPCR analysis relative to *GAPDH* and to controls (n=5 E18.5 embryonic liver samples per genotype). Data shown as mean and S.E.M.; P value calculated by Mann-Whitney t-test. **D**. Analysis of expected and observed numbers of mutant mice before and after birth. P value calculated by a X^2^ test. **E**. Gene Ontology (GO) analysis for biological process comparing embry-onic livers at E18.5 in embryos expressing the fusion protein compared to controls. **F**. Analysis of genes related to hepatocytes and cholan-giocytes in control and transgenic E18.5 livers. Data shown as mean and range. P-adjusted values are shown.

The cell of origin of FLC cannot be determined conclusively from patients. While mouse models have shown that adult hepatocytes can initiate FLC [27, 28], the development of FLC in young patients and the expression of stem cell markers in these tumors [3, 13-17, 44] are also suggestive of an initiating event early in liver development. We wondered if expression of the DNAJB1-PKA fusion in the developing liver may initiate FLC more frequently and more rapidly than in the adult setting. To test this idea, we crossed *Rosa26*^*LSL-DNAJB1-PKA-GFP*^ mice to *Alb-Cre* mice where the Cre recombinase is expressed under the control of the *Albumin* promoter and is turned on at mid-gestation in the liver. Analysis of GFP expression showed positive signal in the liver of mice also expressing Cre at embryonic day E18.5 (**Fig. 1B**). Accordingly, we detected increased expression of *PRKACA* mRNA molecules in transgenic mice expressing Cre at the same time point (**Fig. 1C**).

These observations indicated proper expression of the transgene and led us to investigate the consequences of DNAJB1-PKA expression in the embryonic liver for possible tumor development later in adult mice. However, no *Alb-Cre*;*Rosa26*^*LSL-DNAJB1-PKA-GFP*^ pups from this cross survived past weaning (**Fig. 1D**).

### Expression of DNAJ1B-PKA in the embryonic liver leads to developmental defects

As a first step to understand the cause underlying this perinatal death, we performed RNA sequencing (RNA-seq) from livers dissected from control and transgenic embryos at embryonic day 18.5 (E18.5). The analysis of these data confirmed expression of the fusion transcript in the transgenic livers (**Fig. S1A**,**B**). We identified 331 genes significantly downregulated and 240 genes significantly upregulated in *Alb-Cre Rosa26*^*LSL-DNAJB1-PKA-GFP*^ E18.5 livers compared to controls (**Table S1**). Gene ontology (GO) term analysis showed an enrichment for processes related to inflammation and metabolism, including fatty acid metabolism (**Fig. 1E** and **Table S2**). This analysis was suggestive of differences in liver development. Indeed, when we examined the expression of genes expressed at different stages and in different cell types during liver development [47], we found that a number of genes related to hepatocytes were upregulated in transgenic livers compared to controls. This included *Afp* (coding for Alpha-fetoprotein), *Arg1* (coding for Arginase-1), *Krt18* (coding for cytokeratin 18), as well as genes coding for enzymes implicated in gluconeogenesis (e.g., *Tat* for tyrosine transaminase and *Pck1* coding for phosphoenolpyruvate carboxykinase 1) (**Fig. 1F** and **Table S1**). In contrast, we noted a downregulation of genes coding for markers of cholangiocytes, including *Sox9, Krt7* and *Krt19* coding for cytokeratins 7 and 19, and *Cyp7a1* coding for cholesterol 7 alpha-hydroxylase; there was also upregulation of the neuroendocrine gene *Chgb*, which codes for Chromogranin B; neuroendocrine gene programs have been associated with injured liver phenotypes [48] (**Fig. 1F** and **Table S1**).

These observations indicate that expression of the DNAJB1-PKA fusion in *Albumin*-positive liver cells results in developmental phenotypes that are incompatible with survival.

### DNAJ1B-PKA expression in the adult liver of mice results in the development of slow-growing tumors with low penetrance

Fluorescence in situ hybridization (FISH) shows that the gene fusion is only present in FLC cells and not in surrounding liver cells [49]. To introduce the oncogenic fusion in a subset of adult liver cells and allow for clonal growth, we performed hydrodynamic tail vein tail injections of *Rosa26*^*LSL-DNAJB1-PKA-GFP*^ mice with a plasmid expressing the Cre recombinase (**Fig. 2A**), generating a cohort of 38 animals (28 males and 10 females). We aged these mice and waited for signs of disease before analysis. We found that 10 mice developed tumors (3 spleen tumors, which were not further studied, and 7 liver tumors – **Table S3** and **Fig. 2B**). The low penetrance of FLC development and the slow growth of these tumors in this transgenic model is consistent with previously described mouse models [27, 28], and makes it difficult to use these mice as pre-clinical models of FLC development and response to candidate therapies.

**Figure 2.**
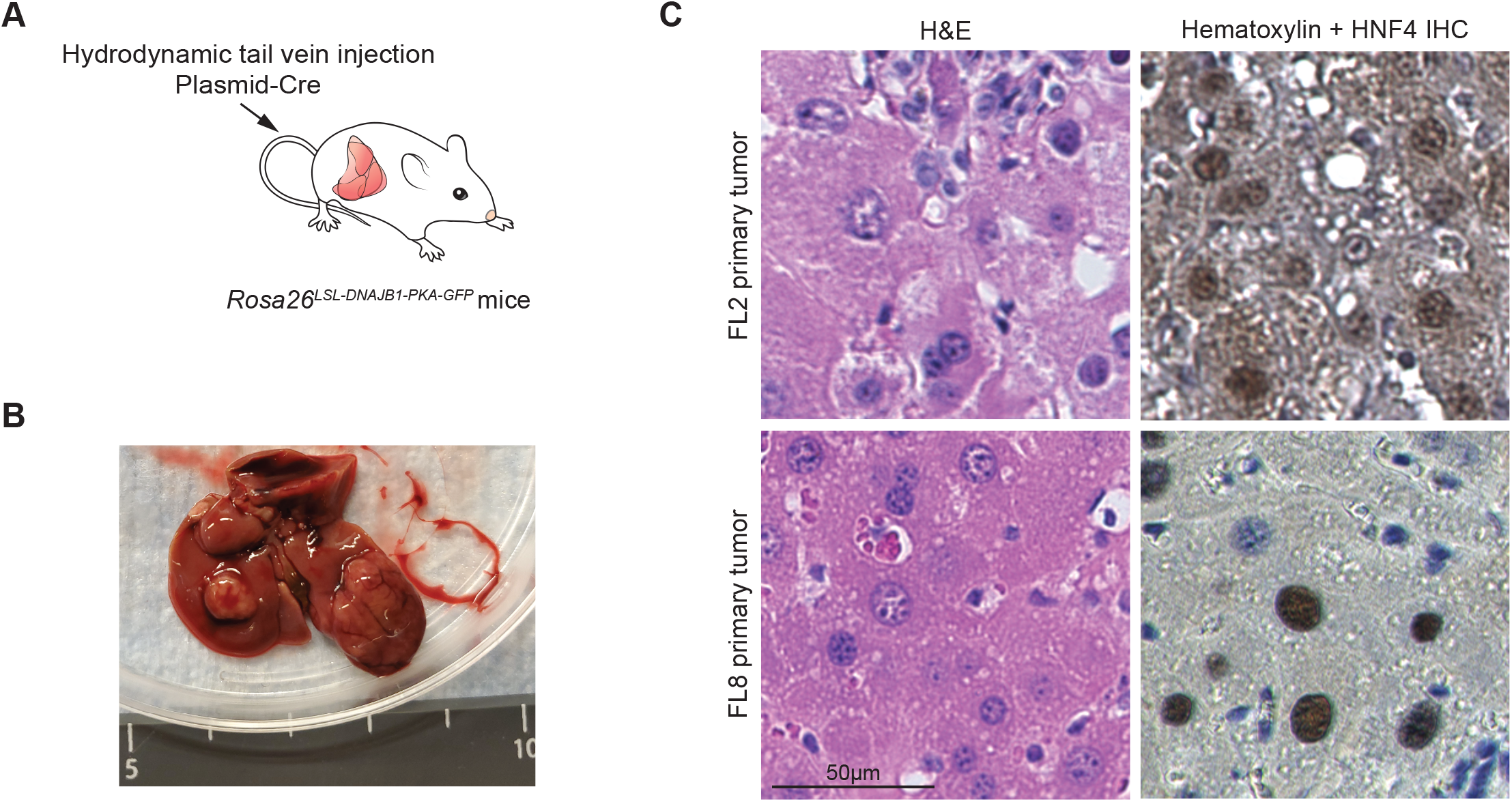
Propagation of tumors from transgenic mice expressing the DNAJ1B-PRAKA fusion. **A**. Schematic representation of the protocol to induce liver tumors in adult *Rosa26*^*LSL-DNAJB1-PKA-GFP*^ mice upon delivery of a plasmid expressing the Cre recombinase to the liver. **B**. Representative photograph of liver tumors in the dissected liver of a *Rosa26*^*LSL-DNAJB1-PKA-GFP*^ mouse 24.5 months after delivery of the Cre recombinase (FL1 tumor shown). **C**. Hematoxylin and eosin and hematoxylin (H&E) staining (left) and anti-HNF4α immunohistochemistry (right) on sections from the FL2 and FL8 primary tumors. Scale bar: 50µm.

We wondered if some of the most aggressive tumors in transgenic mice may be able to be transplanted into recipient mice in allograft models. We were able to generate single-cell suspensions from tumors in 5 out 7 of the mice with liver tumors, 2 of which grew in NSG immunodeficient mice upon subcutaneous injection. One allograft model had a fibrosarcoma histology and was not pursued further (**Fig. S2A**,**B**). The other tumor, FL1 (shown in **Fig. 2B**), had histopathological features of FLC and some GFP expression by immunostaining (**Fig. S2C**,**D**); the first allograft generated from this primary tumor stopped growing when its size was still small and did not further expand in subsequent passages, and it was not further studied.

Based on this low efficiency of tumor take from single cells, we also directly implanted small pieces from 2/7 tumors subcutaneously into NSG mice. These two models, FL2 and FL8, grew and could be propagated into new NSG hosts. Cells in these tumors were positive for the liver marker HNF4α (Hepatocyte nuclear factor 4α) [50] and GFP (**Fig. 2C** and **Fig. S3A**). Similar to the two other mouse models previously described [27, 28], FL2 and FL8 tumors were largely negative for the expression of cytokeratins 17 and 19 (CK17/19) (**Fig. S3B**).

We performed bulk RNA sequencing (RNA-seq) of such passaged tumors. However, when we searched for the transgene mRNA, these cell lines did not express it. In addition, we analyzed the frozen stocks of FL2 and FL8 by genomic DNA PCR and were not able to identify deletion of the stop cassette allowing expression of the transgene (data not shown). Thus, naturally-occurring liver cancer cells likely outcompeted transgene-expressing cells that may have been present in the liver of aged mice.

## DISCUSSION

Here we present a new transgenic mouse model for human FLC based on expression of the DNAJ1B-PKA fusion protein. The *Rosa26*^*LSL-DNAJB1-PKA*^ transgenic model is versatile as Cre can be delivered at different times and in different cell types using other mouse alleles or viral vectors.

When we activated DNAJ1B-PKA during embryonic liver development, mice did not survive to adulthood, suggesting that expression of the fusion protein at this stage is detrimental to some aspect of liver function. Our RNA-seq data suggest that the perinatal lethality may be due to developmental defects, including possibly in bile ducts but future work will be needed to investigate more thoroughly the consequences of DNAJ1B-PKA expression in the embryonic liver. This lethality could be possibly alleviated by decreasing the number of liver cells expressing Cre and may help initiate tumors more efficiently than Cre delivery in the adult liver, for example using an *Alb-CreER* allele with tamoxifen-inducible Cre expression [51], using low doses of tamoxifen during embryogenesis. The *Rosa26*^*LSL-DNAJB1-PKA*^ transgenic model could also be used to model other cancer types in which DNAJ1B-PKA is thought to be oncogenic, including rare cases of pancreatic cancer [52], which may be modeled with the appropriate Cre driver.

A limitation of our model, similar to previous mouse models [27, 28], is the slow and variable development of autochthonous tumors. It is possible that the slow growth of tumors and the development of few tumors is further due to expression of GFP together with the fusion protein, as GFP is immunogenic [53].A growing number of organoid and patient-derived xenograft models are being developed from patients with FLC to investigate key features of this disease (see [29, 30, 45, 46, 54-57] for recent examples). We sought to grow FLC-like cell models in culture and as allografts from our transgenic mouse model. We were able to grow some cell models, but these models did not express the transgene, indicating that the expanding cell populations in this context likely come from the cancerous transformation of normal liver cells with aging (and not from the Cre-mediated activation of the oncogenic fusion). This further illustrates the difficulty of expanding mouse FLC-like cells. Future models may need to include additional genetic alterations to promote more aggressive tumor growth.

It has often been difficult for rare diseases such as FLC to benefit from new therapies because pre-clinical models often lag compared to more frequent tumor types. The development of new mouse models, but also fish models [58], may help provide pre-clinical data that lead to the design of new clinical trials in patients with FLC.

## METHODS

### Mice and generation of tumors

Mice were maintained according to practices prescribed by the NIH at Stanford’s Research Animal Facility (approved protocol #32398). Additional accreditation of Stanford animal research facilities was provided by the Association for Assessment and Accreditation of Laboratory Animal Care (AAALAC).

A *Rosa26*^*LSL-DNAJB1-PKA-GFP*^ C56BL/6 founder mouse was generated and validated by Applied StemCell using a site-specific integrase via pronuclear injection [59]. *Alb-Cre* mice were described before [60] and obtained from the Jackson Laboratory (Stock No: 035593). Primers for genotyping are available upon request.

To generate tumors in the liver of mice, 10-16-week-old mice were anesthetized using isoflurane, the tail vein was dilated with a heat pad, and 10 µg of Turbo-Cre plasmid was delivered in a solution of sterile PBS using a hydrodynamic tail-vein injection protocol. The Turbo-Cre plasmid was a gift from the laboratory of Dr. Steven Artandi at Stanford University. Mice were housed for 1-2 years before subsequent analysis.

### RT-qPCR

Liver tissue was isolated by microdissection, minced with a razor blade, and total RNA was collected using RNeasy Fibrous Tissue Mini Kit (Qiagen). cDNA synthesis was performed using the iScript cDNA synthesis kit (Bio-Rad) and RT-qPCR was performed using iQ SYBRGreen supermix (Bio-Rad). The following RT-qPCR primers were used DNAJPRKACA F: 5’-TTACTACCAGACGTTGGGCCT-3’, DNAJPRKACA R: 5’-ATAGTGGTTCCCGGTCTCCT-3’, mPRKACA-aF: 5’-AGATCGTCCTGACCTTTGAGT-3’, mPRKACA-aR: 5’-GGCAAAACCGAAGTCTGTCAC-3’, mPRKACA-bF: 5’-GGTGACAGACTTCGGTTTTGC-3’, mPRKACA-bR: 5’-CACAGCCTTGTTGTAGCCTTT-3’.

### Immunostaining

Tissues were fixed with 4% paraformaldehyde (PFA) in phosphate-buffered saline (PBS). Liver tissues (normal or tumor) were initially perfused with 10 mL PFA through the inferior vena cava. All tissues were submerged in 4% PFA for 48 hr before being switched to 70% ethanol and finally processed and embedded. Slides were deparaffinized in 2x 5 min changes of Histo-Clear (National Diagnostics HS-202), and rehydrated in 2x 1 min changes of 100% ethanol, 2x 1 min changes of 70% ethanol, and finally 2x 5 min changes of distilled water. Antigen retrieval was performed in a microwave for 10 min with citrate-based buffer (Vector Labs H-3300). Slides were cooled at room temperature, quenched for endogenous enzyme activity (SP-6000), and washed in 3x 3 min changes of PBS with 0.1% Tween-20 (PBST) (Sigma P1379). Slides were blocked with secondary antibody serum for 1 hr at room temperature (RT). The GFP antibody (Cell Signaling Technology #2956) was diluted to 1:200 in PBST and applied to slides for 1 hr at 37 ºC. Slides were washed in 3x 3 min changes of PBST. Horse anti-rabbit secondary (Vector Labs MP-7401) was applied to slides for 1 hr at 37 ºC. Slides were washed in 3x 3 min changes of PBST. The chromogen was developed using a DAB substrate kit (Vector SK-4100) for 5 min before quenching in distilled water for 5 min. Slides were counterstained with hematoxylin (Newcomer Supply 1201B, dehydrated in 2x 1 min changes of 70% ethanol, 2x 1 min changes of 100% ethanol, and finally cleared in 2x 5 min changes of xylene before mounting.

Embryos were dissected from timed pregnant females, fixed in 4% PFA overnight and sections were generated as described above. Sections were depaffarinized, rehydrated, and tissue was blocked in 5% goat serum for 1 hr. Anti-GFP was added overnight. The following day, samples were washed before incubation with Goat anti-Rabbit - Alexa Fluor™ 488 (Thermo Fisher Scientific) and DAPI (Sigma). Samples were washed and mounted in ProLong Gold mounting media (Thermo Fisher Scientific) before confocal imaging.

### Tumor transplantation studies

Nod.Cg-Prkdc^scid^IL2rg^tm1WjI^/SzJ (NSG) mice (Jackson Laboratories, Stock No: 005557) were used for all experiments in immunodeficient recipients.

Tumors from *Rosa26*^*LSL-DNAJB1-PKA-GFP*^ mice were microdissected from livers or lungs. Tumors were bisected once to increase surface area exposed to transplant media (Kubota’s medium) and kept on ice until ready for transplant. Recipient mice were anesthetized. The shaved flank was sterilized with one swab of betadine, one swab of 70% ethanol repeated 3 times. After confirming adequate depth of anesthesia by toe-pinch, a 5 mm incision was made in the shaved flank. Blunt dissection under the skin was used to separate the skin from the muscle, creating a pocket between the 2 layers. Tumor pieces were implanted (bisected side facing the muscular wall, ∼5 mm piece) near macroscopically visible capillaries on muscle wall. The incision was closed using 9 mm wound clips. For orthotopic injections, cells were disaggregated from Matrigel using the protocol described above. Cell pellets were resuspended at a concentration of 10^4^-10^5^ cells in 25 µL. These cell suspensions were mixed with an equivalent volume of Matrigel to a total volume of 50 µL. Mice were anesthetized and shaved in the abdomen. Lateral incision was made in the skin directly inferior to xyphoid process. Blunt dissection was used to separate surrounding skin from parenchyma and musculature. Incision was made in caudal direction originating from xyphoid process. The largest liver lobe available was exposed and stabilized. For cell injection, 20 µL of cells mixed 1:1 with Matrigel were injected. For tissue implantation, tissue 3-8 mm-wide at longest axis was sutured to the liver lobe using 5-0 vicryl suture. Liver lobe was returned to original site. Mouse musculature was closed with 5-0 vicryl running suture, skin was closed with 9mm wound clips.

### RNA sequencing analysis

Frozen tumor samples were sent to Novogene for RNA extraction and library preparation. Raw sequencing reads were demultiplexed by the vendor to generate approximately 20 million paired-end reads per sample. Fastq files were trimmed using CutAdapt (v2.10) [61] using TruSeq sequencing adapter (5’-AGATCGGAAGAGCACACGTCTGAACTCCAGTCAC-3’), and a minimum read length of at least 25. Reads were then aligned to the ENSEMBL mm10 genome using HiSat2 [62] using reverse strandedness and discarding unaligned reads. Counts were assigned to genes using featureCounts [63] at ENSEMBL gene annotation (v93). Differential expression analysis was conducted using DESeq2 [64] on R v4.1.2 (https://www.R-project.org/).

### Statistics and reproducibility

Statistical significance was assessed using the Prism GraphPad software. The specific tests used are indicated in the figure legends. Investigators were not blinded to allocation during experiments and outcome assessment.

## Data availability

All RNA sequencing datasets generated in this study are available at Gene Expression Omnibus (GEO) under Super Series GSE249325 (https://www.ncbi.nlm.nih.gov/geo/query/acc.cgi?acc=GSE249325). All other data are available in the article and supplementary materials, or from the corresponding author upon reasonable request.

## ACKNOWLEDGMENTS

We thank Pauline Chu from the histology facility for her help with tissue sections and all the members of the Sage lab for their help and support throughout this study. Research reported in this publication was supported by grants from the Fibrolamellar Cancer Foundation and the Alex’s Lemonade Stand Foundation. J.S. is the Elaine and John Chambers Professor in Pediatric Cancer.

## Notes

https://www.ncbi.nlm.nih.gov/geo/query/acc.cgi?acc=GSE249325

### Competing Interest Statement

The authors have declared no competing interest.

### Summary of Updates

We added new data (RNA sequencing analysis) to characterize the embryonic liver phenotypes in the transgenic mice expressing the oncogenic fusion. In addition, the first version of this manuscript described established cell models from mouse tumors but, upon genotyping these cell models, we realized the growing cancer cells were contaminating liver cancer cells not driven by the oncogenic fusion (naturally-occurring tumors in aging animals). We thus removed this part from the manuscript.

