## Supplemental Figures S1-3 for "Development of pre-clinical murine models for fibrolamellar hepatocellular carcinoma"

### FIGURE S1

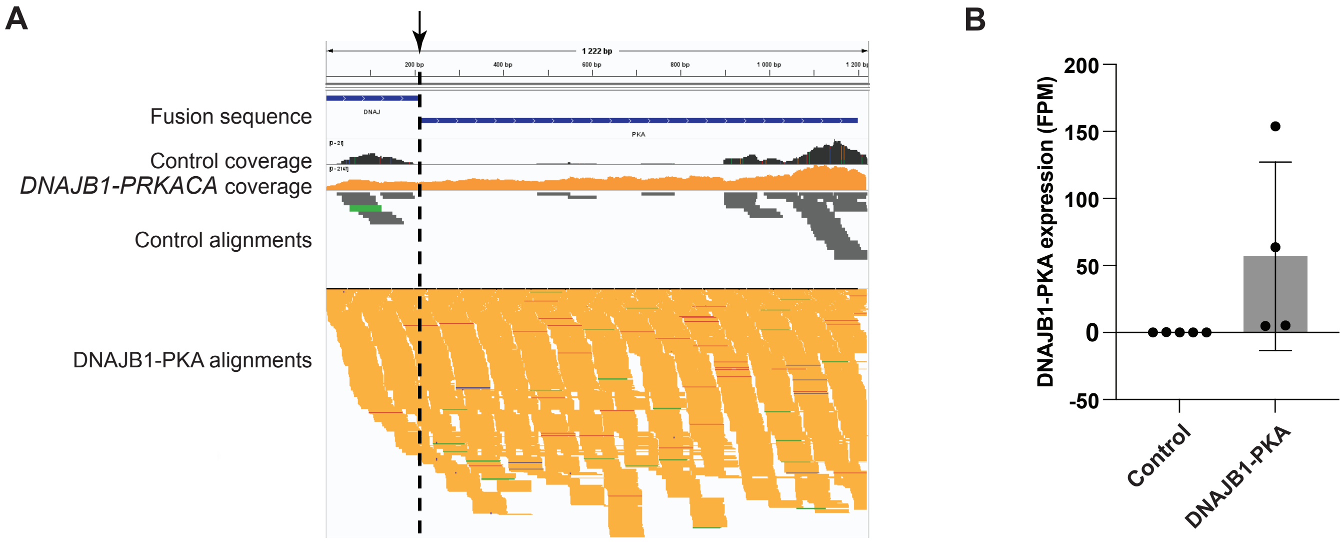


**Figure S1. RNA-seq analysis of control and transgenic embryos expression the fusion transcript**

**A.** Analysis of *DNAJB1-PRKACA* fusion transcript reads from the RNA-seq analysis. **B**. Expression levels of the *DNAJB1-PRKACA* fusion transcript from the RNA-seq analysis (adjusted p-value <2.7e-8 from a DEseq analysis).

### FIGURE S2

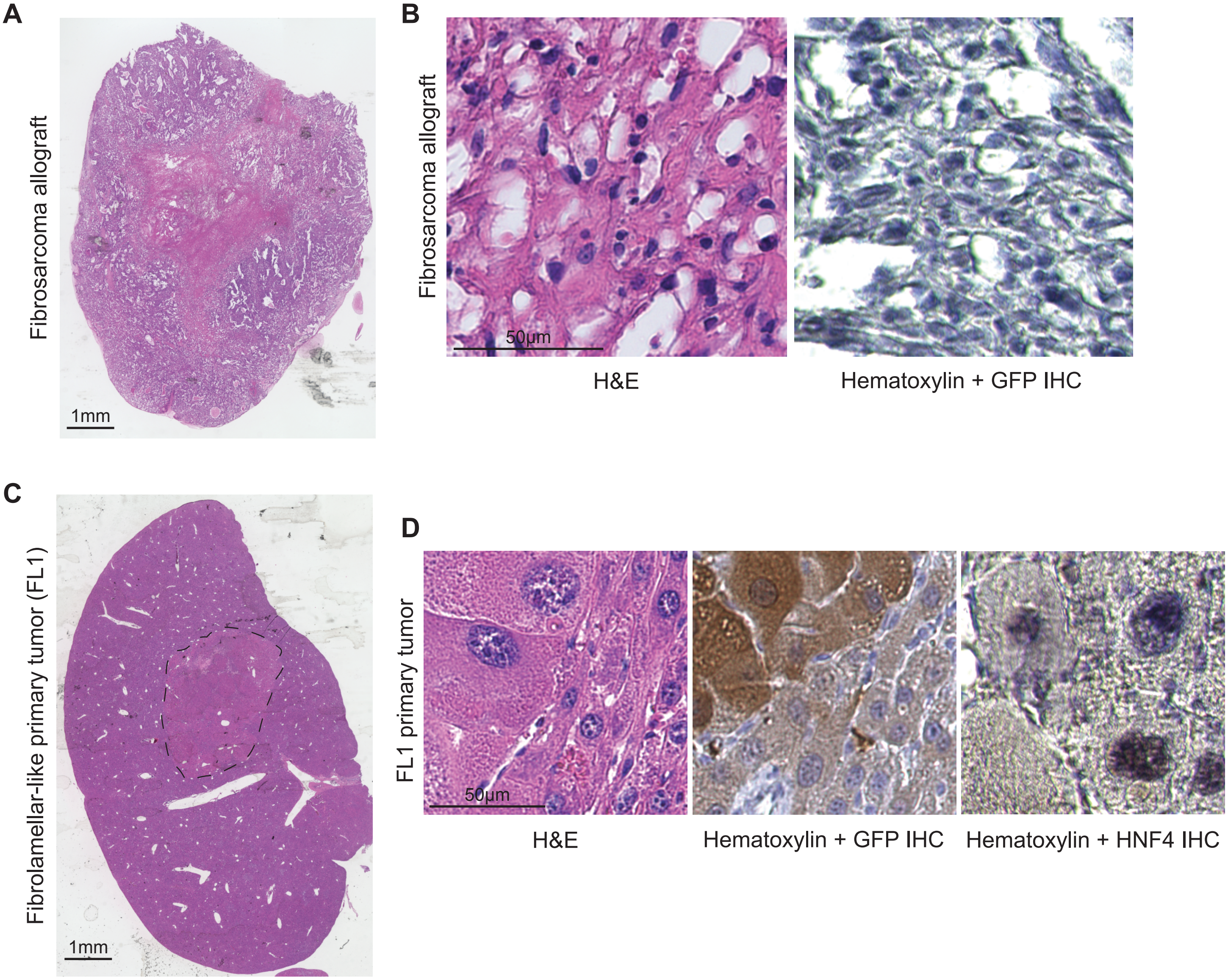


**Figure S2. Tumors growing in the liver of adult *Rosa26*$^{LSL\text{-}DNAJB1\text{-}PKA\text{-}GFP}$ mice**

**A.** Hematoxylin and eosin (H&E) staining of a fibrosarcoma-like liver tumor growing in the flank of an NSG host, at low magnification. Scale bar: 1mm. No tumor sample was left from the original liver tumor. **B.** Higher magnification view of the allograft in (A), with GFP immunohisto-chemistry; note the absence of brown signal that would be indicative of GFP expression. Scale bar: 50µm. **C.** Hematoxylin and eosin (H&E) staining of a fibrolamellar-like liver tumor (dashed line) (FL1 model) at low magnification. Scale bar: 1mm. **D.** Higher magnification view of the primary tumor in (C), with GFP (transgene) and HFN4α (liver origin) immunohistochemistry (brown signal). Scale bar: 50µm.

#### FIGURE S3

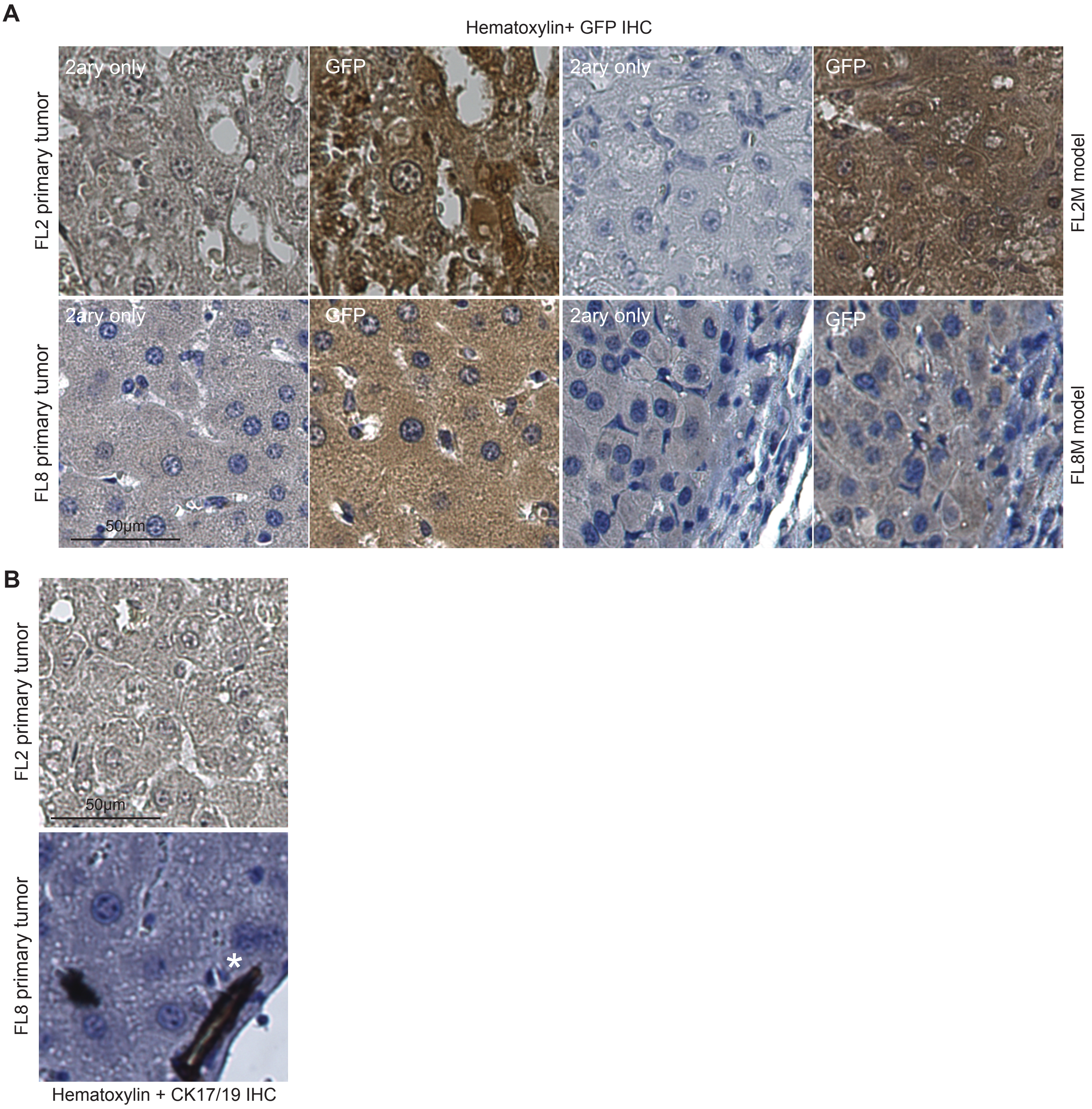


**Figure S3. Propagation of fibrolamellar-like tumors and metastases from** ***Rosa26***$^{LSL\text{-}DNAJB1\text{-}PKA\text{-}GFP}$ **mice**

**A.** Hematoxylin staining of sections from the four models indicated together with anti-GFP immunohistochemistry, with the secondary antibody-only control. Scale bar: 50µm. **B.** Hematoxylin staining of sections from the two tumors indicated together with anti-CK17/19 immunohistochemistry. The asterisk shows a bile duct (brown positive signal) as a positive control. Scale bar: 50µm.
